# Presence of *Legionella* spp. in cooling towers: the role of microbial diversity, *Pseudomonas*, and continuous chlorine application

**DOI:** 10.1101/540302

**Authors:** Kiran Paranjape, Émilie Bédard, Lyle G. Whyte, Jennifer Ronholm, Michèle Prévost, Sébastien P. Faucher

## Abstract

Legionnaire’s Disease (LD) is a severe pneumonia caused by *Legionella pneumophila*. Cooling towers are the main source of *L. pneumophila* during large outbreaks. Colonization, survival, and proliferation of *L. pneumophila* in cooling towers are necessary for outbreaks to occur. These steps are affected by chemical and physical parameters of the cooling tower environment. We hypothesize that the bacterial community residing in the cooling tower could also affect the presence of *L. pneumophila*. A *16S rRNA* targeted amplicon sequencing approach was used to study the bacterial community of cooling towers and its relationship with the *Legionella spp.* and *L. pneumophila* communities. The results indicated that the water source shaped the bacterial community of cooling towers. Several taxa were enriched and positively correlated with *Legionella spp.* and *L. pneumophila*. In contrast, *Pseudomonas* showed a strong negative correlation with *Legionella spp.* and several other genera. Most importantly, continuous chlorine application reduced microbial diversity and promoted the presence of *Pseudomonas* creating a non-permissive environment for *Legionella spp*. This suggests that disinfection strategies as well as the resident microbial population influences the ability of *Legionella spp.* to colonize cooling towers.

## INTRODUCTION

Legionnaires’ Disease (LD) is a severe and potentially fatal pneumonia caused by several bacterial species of the genus *Legionella*. More than 90% of cases are caused by the species *Legionella pneumophila* [1]. The remaining 10% of cases are caused by other species, such as *L. longbeachae*, *L. bozemanii*, and *L. dumoffii* [2, 3, 4]. LD is usually contracted through the inhalation of contaminated aerosols. Consequently, Engineered Water Systems (EWS), such as hot water distribution systems, cooling towers, water fountains, misters, and whirlpool spas are sources of dissemination of the bacterium [5, 6, 7, 8, 9, 10]. Cooling towers are the major source for large outbreaks and up to 28% of all sporadic cases [11, 12].

In recent years, the number of cases of LD has increased both in Europe and North America [13, 14]. From 2000 to 2014, the CDC reported an increase of 286% in cases of Legionellosis (LD and Pontiac fever) in the USA [15]. This increase is likely due to increasing population in urban areas, improvements in surveillance methods, aging populations, and climate change [13]. *Legionella* is now the main cause of death due to waterborne diseases in the US [16].

Several steps are needed for a tower to become the source of an outbreak of LD. First, the tower must be seeded with *L. pneumophila*. During operation, the water lost through evaporation is replenished either with municipal water, onsite well water or available surface water, which may be the source of *L. pneumophila* [17, 18]. Next, *L. pneumophila* must survive and proliferate in the cooling tower environment. Encountered stresses include low quantity of nutrients, disinfectants, and competing microbes [19, 20]. *L. pneumophila* can survive up to several months in oligotrophic water while retaining infectivity [21]. Multiple factors may affect the prevalence of *Legionella* and its hosts in cooling towers including operational factors, temperature, water quality, the age of the equipment, the use of biocides (dosage, type and application schedule and residual concentration), and elevated bacterial indicators such as heterotrophic plate counts (HPC) [22, 23, 24, 25]. In addition, biofilms offer protection against disinfectants, while also providing nutrients and host cells [26, 27, 28, 29, 19]. While it is not clear if *L. pneumophila* can grow in biofilms independently of protozoan host cells, several studies indicate that this might be possible [30, 28, 31]. Moreover, the ability of *L. pneumophila* to colonize biofilms may depend on the microbial community composition of these biofilms [27, 32]. For example, *L. pneumophila* persists in *Klebsiella pneumoniae* biofilms but not in *Pseudomonas aeruginosa* biofilms [27]. In addition, the surface material on which biofilms grow seemed to influence *L. pneumophila* survivability [33, 34, 35, 36]. Finally, some microorganisms present in cooling towers can prey on *L. pneumophila* and reduce its population. For instance, protozoa, such as *Solumitrus palustris*, and bacteria, such as *Bdellovibrio spp.*, feed on *L. pneumophila* in experimental settings [37, 38, 39]. Consequently, the presence of these species may restrict *L. pneumophila*’s colonization of cooling towers.

Following this initial colonisation, the *L. pneumophila* population must grow to sufficient number to be dispersed effectively and cause an LD outbreak. *L. pneumophila* is an intracellular parasite of amoeba and ciliates, such as *Acanthamoeba castellanii*, *Vermamoeba vermiformis* and *Tetrahymena pyriformis* [40, 41, 42]. Consequently, the cooling tower must harbor a large number of host cells in order for *L. pneumophila* to grow sufficiently to contaminate the aerosols produced. The host cell population is also affected by the chemical and physical parameters of the cooling tower environment [43, 44, 45]. As these host cells graze on the bacterial community of cooling towers, microbial interactions necessarily impact their growth. For instance, some host cells may require specific prey in order to grow [37]. Conversely, certain species of bacteria are able to resist predation and even grow intracellularly, effectively competing against *L. pneumophila* [46]. In contrast, *Fischerella spp.* (*Cyanobacteria*) and *Flavobacterium* promote the growth of *L. pneumophila*, [47, 48].

The majority of cooling towers seem to contain a core *Legionella spp.* community [49, 11, 50]. However, the stability of this community is still not well understood and *L. pneumophila* seems able to proliferate to the detriment of other *Legionella* species [49, 50]. Chemical disinfection is a disruptor to the *Legionella* community but *L. pneumophila* seems quicker to recover after chlorine treatment and can dominate the *Legionella* community [49, 50]. Moreover, relative abundance of the family *Legionellaceae* is positively correlated with alpha diversity [11], suggesting that microbial interactions are essential for the growth of *L. pneumophila* in cooling towers.

Thus, outbreaks of LD are driven by chemical and physical properties, as well as microbial interactions. Nevertheless, the ecology of *L. pneumophila* in cooling towers is still poorly understood and potential interactions with resident microbes need to be clarified. Consequently, we used a *16S rRNA* targeted amplicon sequencing approach to characterize the bacterial community of cooling towers, along with the chemical and physical characteristics, and investigate their relationship with *L. pneumophila*. We hypothesize that the presence of *L. pneumophila* depends on certain groups of bacteria, whose presence is influenced by other factors such as disinfectant or water characteristics.

## MATERIALS AND METHODS

### Sampling of cooling towers

A total of 18 cooling towers were sampled from six different regions in Quebec, Canada, between the 10^th^ and 21^st^ of July 2017. Location of towers, total and residual chlorine levels, and disinfection regimes are listed in Table 1. Water was sampled with sterile polypropylene bottle from the basin of the cooling tower or from a sampling port when the basin was inaccessible. All towers were sampled in triplicate in volumes of one litre to perform heterotrophic plate counts and *16S rRNA* targeted amplicon sequencing. An additional two litres were collected to analyze chemical and physical parameters. Samples were brought back to the lab stored at room temperature and processed within 48 hours.

**Table 1:**
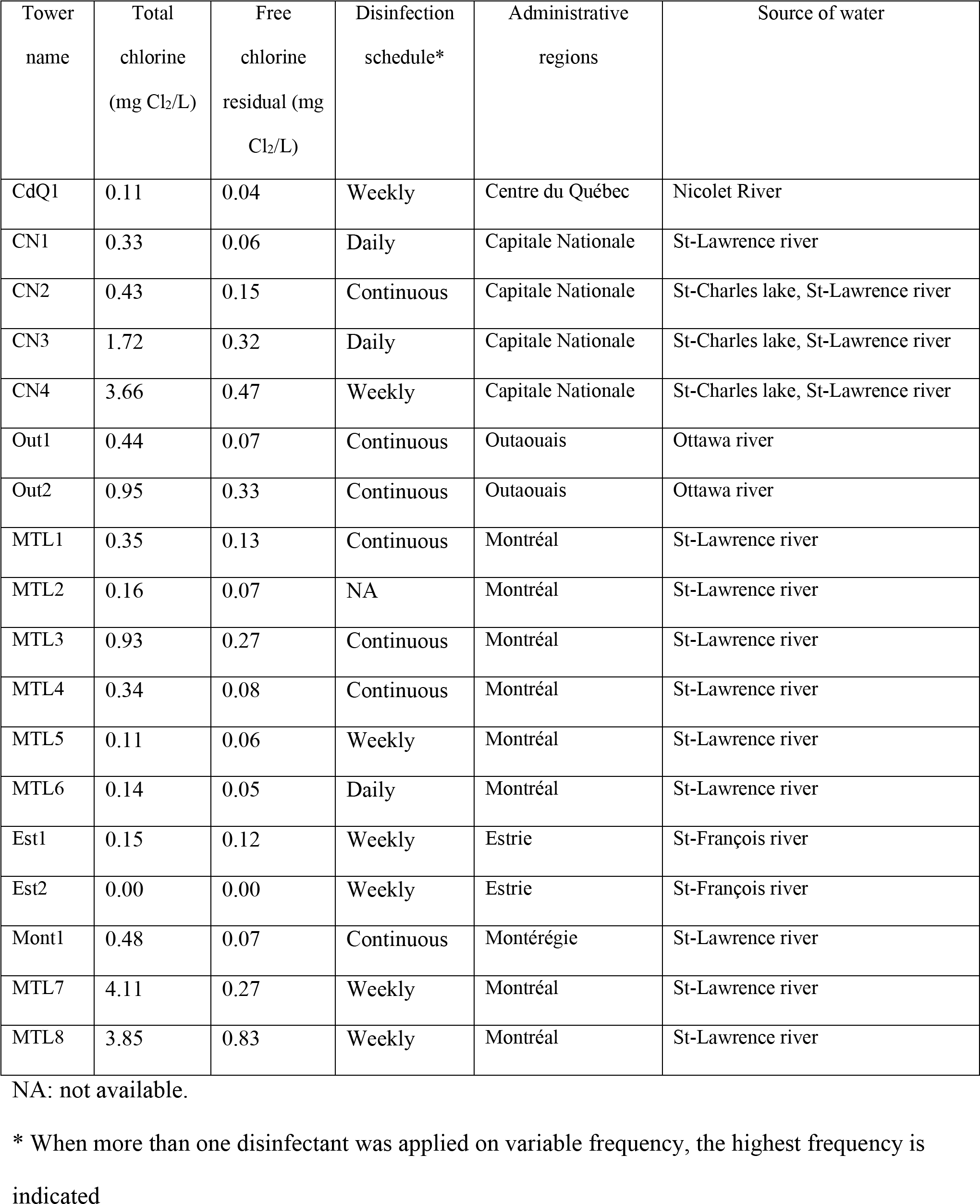
Disinfection program and Location of cooling towers.

### Heterotrophic plate count, physical and chemical parameter measurements

Heterotrophic plate count (HPC) were performed on R2A and nutrient agar media, which were incubated at 30°C for 24 hours. Turbidity, pH, temperature, total chlorine, residual chlorine, conductivity and dissolved oxygen were measured on-site. Residual and total chlorine were measured using a Pocket Colorimeter™ II (Hach, Loveland, CO, USA), conductivity, turbidity with a Hach 2100Q (Hach, Loveland, CO, USA) while pH and dissolved oxygen were measured using a Hach Multi-Parameter HQ40d tool (Hach, Loveland, CO, USA). Water samples were further analysed for the following chemical parameters: total suspended solids (TSS) and suspended volatile solids (VSS, Standard Methods 2540D, E), dissolved organic carbon (DOC, Standard Methods 5310C with 0.45 um filtration), biodegradable dissolved organic carbon [51], dissolved and total iron (Inductively Coupled Plasma). Nitrite, nitrate, ammonia, phosphorus, sulphide, and sulphate were measured using colorimetric kits (CHEMetrics, Midland, VA, USA) according to the manufacturer’s instruction.

### Filtration of biomass and DNA extraction

Water samples were filtered through 0.45 μm pore size mixed cellulose ester membrane filters (Millipore, Burlington, MA, USA). Each replicate was filtered and processed separately. The DNeasy PowerWater Kit from Qiagen (Cat. No. 14900-100-NF, Germantown, MD, USA) was used to extract DNA from the filters. The manufacturer’s protocol was followed, except that nuclease-free water was used for the final elution step. The extracted DNA was quantified using a Nanodrop (Thermofisher, MA, USA) and the purified DNA was stored at −20°C.

### Bacterial profiling of cooling towers using 16S rRNA targeted amplicon sequencing

*16S rRNA* targeted amplicon sequencing was performed on the Illumina MiSeq platform (Illumina, inc) using a sequencing strategy developed by *Kozich et al*, which uses a dual index sequencing strategy using the F548 and R806 primers which amplify the V4 region of the bacterial *16S rRNA* gene [52]. Briefly, the V4 region of the bacterial *16S rRNA* was amplified using the Hot Start Taq Plus Master Mix (Qiagen, Germantown, MD, USA) and indexed primers [52]. The cycling program consisted of an initial denaturation step of 95°C for 2min, followed by 25 cycles of 95°C for 20 seconds, 55°C for 15 seconds, and 72°C for 5 minutes, and a final elongation of 10 minutes at 72°C. The PCR products were then purified using Ampure XP beads (Beckman Coulter, Indianapolis, IN, USA) according to the manufacturer’s instruction. The purified DNA was quantified using the Quant-iT PicoGreen dsDNA assay kit (Thermofisher, MA, USA). The DNA samples were then normalized to a concentration of 1.5 ng/μl, pooled together, mixed with 10% PhiX sequencing control (Illumina, inc), diluted to 4.0 pM, and denatured with a final concentration of 0.0002N NaOH. The sequencing run was performed on the MiSeq platform with the MiSeq Reagent kit V2, according to the manufacturer’s instruction. Raw sequence reads were deposited in Sequence Read Archive under accession number PRJNA507738.

Sequencing data was processed using the Mothur pipeline [52]. Briefly, the paired reads were assembled into contigs, and any contig with ambiguous bases or longer than 275bp were culled. Sequences were aligned to the bacterial Silva reference database release 132. Sequences that did not align to the reference database were removed. The ends and gaps from the sequence alignment were trimmed so that all sequences had the same alignment coordinates. The sequences were further denoised using a pre-cluster algorithm implemented in Mothur. The resulting unique sequences were purged of chimeras using the VSEARCH algorithm. Additionally, any undesirable sequences remaining, such as Eukaryota, Archaea, chloroplasts, and mitochondria, were removed using a Bayesian classifier algorithm in Mothur. Next, the sequences were grouped according to their taxonomy and clustered into OTUs at 97% similarity. The MicrobiomeAnalyst web-based tool was used to analyse the OTU data and perform LEfSe analysis (http://www.microbiomeanalyst.ca/faces/home.xhtml) [53]. OTUs with low counts were filtered out using the default parameters (at least 20% of the samples contain 2 counts or more). One of the replicates for tower CN4 had significantly lower read levels than all the other samples. Thus, this replicate was omitted from the analysis, and the remaining samples were rarefied to the next lowest read count sample (20 712 sequences). Only duplicates were analysed for tower CN4. GraphPad Prism 7.03 was used to produce most of the graphs along with some statistical analysis.

### Quantification of *L. pneumophila*

*L. pneumophila* was quantified from the DNA extract using the iQ-check *L. pneumophila* quantification kit (Bio-Rad), according to the manufacturer’s instruction. The qPCR was run with a BioRad CFX Connect Real Time system thermocycler. The data was analyzed with the CFX manager 3.1 and GraphPad Prism 7.03. The results are expressed as genome unit per litre (GU/L).

## RESULTS

### Characteristics of cooling towers included in this study

Eighteen cooling towers were sampled between the 10^th^ and 21^st^ of July 2017. Characteristics of each cooling tower as well as water profiles are described in Table 1 and Supplementary Table S1. On average, the water of cooling towers sampled had the following characteristics: temperature, 25.2 ± 2.4 °C; pH, 8.7 ± 0.2; conductivity, 881 ± 275 μS/cm; dissolved oxygen, 8.0 ± 0.5 mg/L; dissolved organic carbon, 17 ± 10 mg/L. As seen in Figure 1, HPC were highly variable, ranging from 10^5^ CFU/L for tower MTL3 to 10^9^ CFU/L for tower Out1. Only five towers (CdQ1, CN2, CN3, MTL5 and Est2) showed detectable level of *L. pneumophila* ranging from 300 to 1300 GU/L (Figure 1), below the regulatory standards [54]. Of note, *L. pneumophila-* positive towers were not restricted to a particular region.

**Figure 1:**
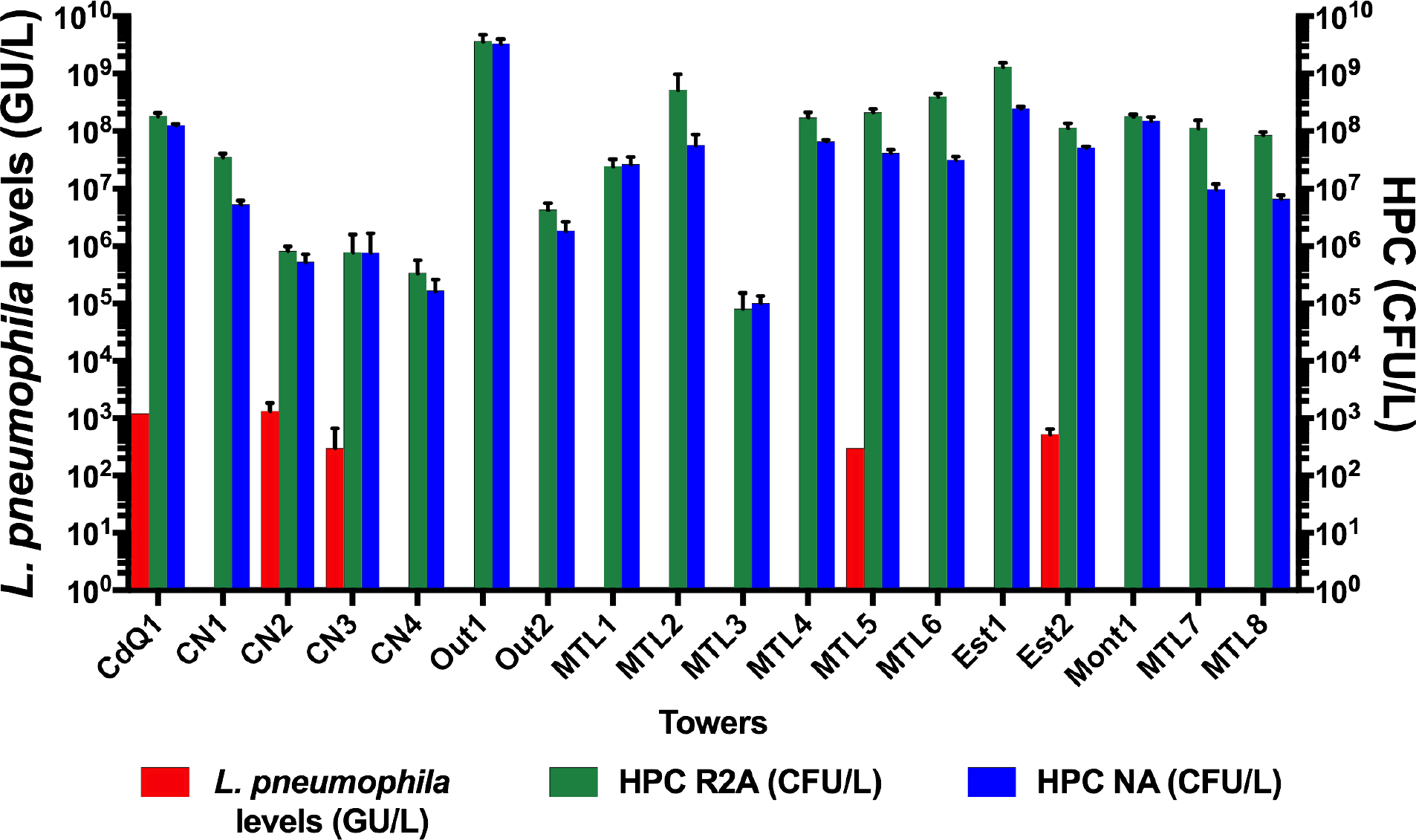
Levels of *L. pneumophila* in genomic units per litre (GU/L) detected by qPCR, and HPC measured on R2A and nutrient agar (NA). The data presented are the average and standard deviation of three sampling replicate. See table 1 for tower name and location details.

### Characterisation of the bacterial community of cooling towers

*16S rRNA* targeted amplicon sequencing was performed on sampling triplicates to study the bacterial makeup of the cooling towers. *Proteobacteria* dominated the bacterial population of all towers at the phylum level (Figure 2A). *Cyanobacteria* and *Bacteroidetes* were the second and third most abundant phyla. Seven towers showed a *Cyanobacteria* population above 1%, which in some cases reached up to around 30% of the entire population (tower CN3). In all cases, the *Cyanobacteria* population consisted of non-photosynthetic candidate phylum *Melainabacteria* [55]. In the case of *Bacteroidetes*, only five towers had a population above 1% reaching 10% for towers CdQ1 and MTL6. On average, the other phyla, such as *Actinobacteria* or *Firmicute*, constituted less than 1% of the population.

**Figure 2:**
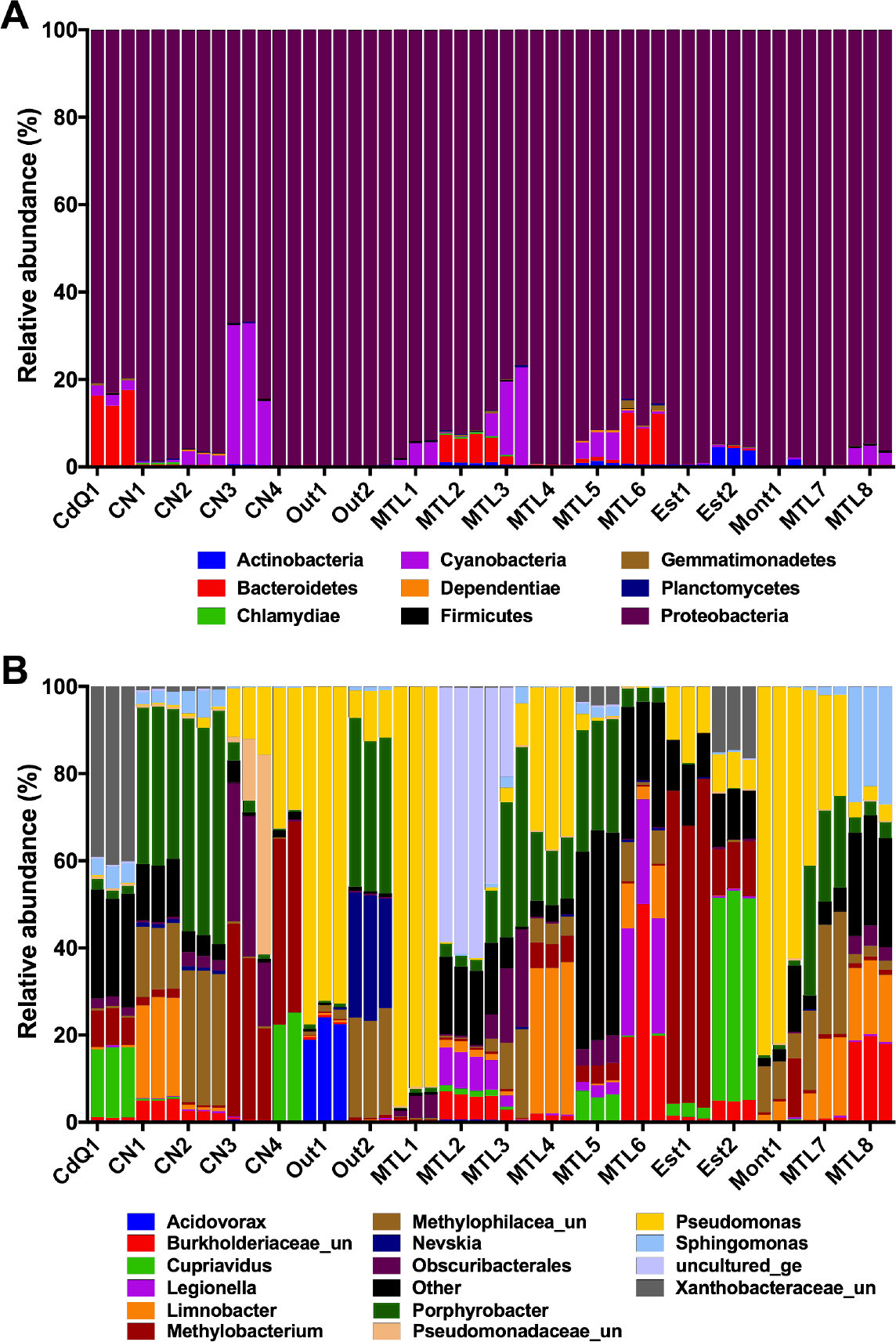
Relative abundance of bacterial OTU classified at the phylum level (A) and at the genus level (B) of the different cooling towers sampled in Quebec, Canada during the summer of 2017. See table 1 for name and location details.

The bacterial populations were also examined at lower taxonomic levels (Figure 2B). Overall, a total of 72 genera passed the low count filter described in the materials and methods section. The relative abundance patterns were similar between replicates but varied greatly between towers (Figure 2B). Several genera commonly found in other water systems were identified, such as *Pseudomonas*, *Limnobacter*, *Porphyrobacter*, *Legionella*, *Cupriviadus* and *Mycobacterium*. Interestingly, rare and uncharacterized genera were also identified, such as *Yonghaparkia* and Tra3-20 [56, 57, 58]. Methylotrophs were found in all towers, with groups such as *Methylobacterium* or unclassified *Methylophilaceae* being highly abundant in some. For instance, more then 70% of the bacterial population of tower Est1 belonged to the *Methylobacterium* genus.

### Effect of water chemistry on alpha diversity of cooling towers

The Shannon diversity index was used to measure alpha diversity. The average Shannon index varied significantly from tower to tower (Kruskal-Wallis, *P* < 0.0001; *H*=47.612; Supplementary Figure S1). TSS, VSS, DOC, total iron, and dissolved iron negatively affected alpha diversity (Supplementary Figure S2). High conductivity was associated with higher alpha diversity (Supplementary Figure S2). Next, the effect of chlorine concentration on alpha diversity was investigated. A threshold of 0.3 mg Cl_2_/L was used to categorize the towers into low and high chlorine groups. Measured total and residual chlorine had no effect on alpha diversity (Figure 3A and B). The frequency of application of chlorine had a significant effect on alpha diversity: continuous chlorination reduced alpha diversity compared to periodic application (daily and weekly, *P* < 0.004, Figure 3C). This suggests that the frequency of application of chlorine has a stronger impact on the microbial diversity of cooling towers than the concentrations of chlorine at the time of sampling.

**Figure 3:**
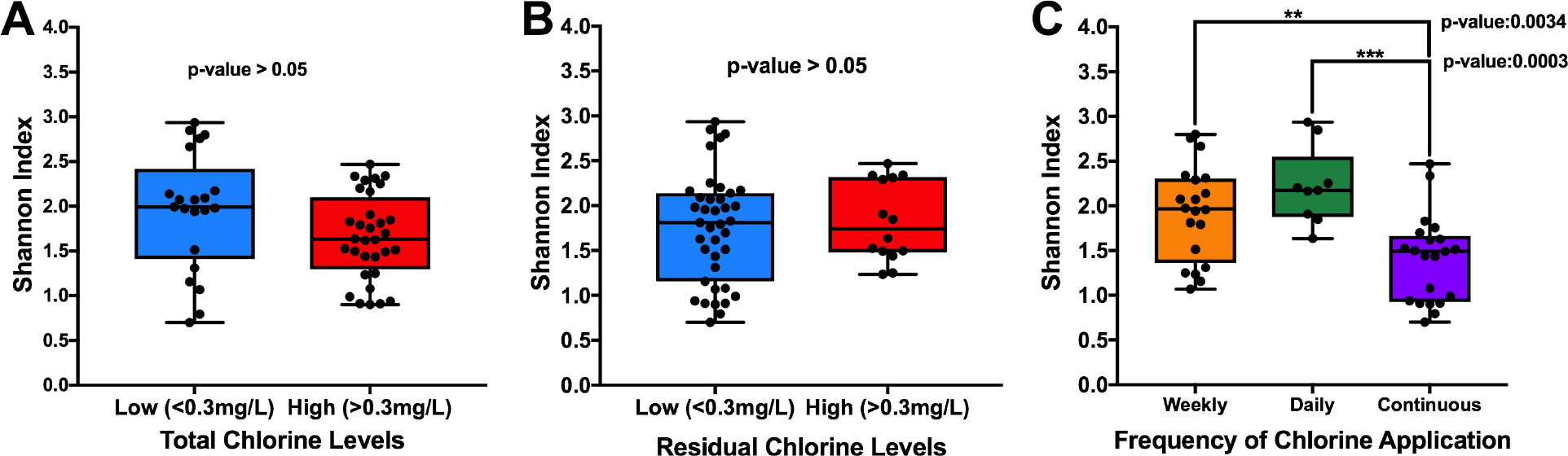
Alpha diversity of cooling towers categorized by levels of total chlorine (A), free residual chlorine (B), and frequency of application (C). In A and B, a Mann-Whitney test was used to assess statistical significance. In C, a Kruskall-Wallis test followed by Dunn’s test for pairwise comparison of samples was used to test statistical significance.

Finally, the effect of alpha diversity on *Legionella*, *Mycobacterium* and *Pseudomonas* was investigated. Some members of *Mycobacterium* and *Pseudomonas* are opportunistic pathogen associated with EWS. *Legionella* and *Mycobacterium* were not present in all samples and, consequently, samples were partitioned into samples containing or not containing these genera. The mean Shannon index for samples without *Legionella* was 1.04, whereas the index was 1.92 for samples with *Legionella* (*P* < 0.0001, Figure 4A). This positive correlation was previously reported for cooling towers located in the United States [11]. The same relationship was observed for *Mycobacterium* (Supplementary Figure S3). No significant differences in alpha diversity were observed between *L. pneumophila-*positive towers and negative towers (*P* > 0.05, Figure 4B), indicating that alpha diversity is not correlated with *L. pneumophila*; however, the low number of positive towers could hide a relationship. Finally, the relation between *Pseudomonas* and alpha diversity was investigated by plotting the *Pseudomonas* reads of each tower against their respective Shannon index. The data followed a non-linear regression model and indicated that alpha diversity of the towers decreased exponentially as *Pseudomonas* read counts increased (R^2^ = 0.78, Figure 4C).

**Figure 4:**
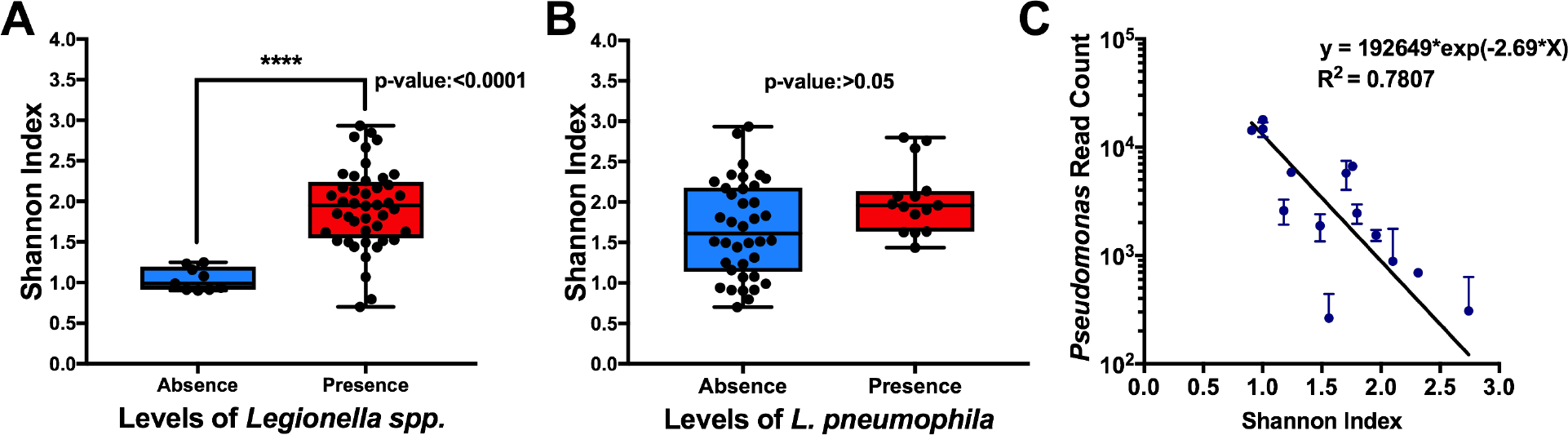
Relationship between alpha diversity and *Legionella* and *Pseudomonas*. The cooling towers categorized by (A) the presence of *Legionella spp.* according to 16S rRNA amplicon sequencing and (B) *L. pneumophila* detected by qPCR. The Mann-Whitney test was used to determine statistical significance. (C) *Pseudomonas* reads were plotted against the Shannon index of each tower.

### Effect of geographic location on the microbiome

Beta diversity was calculated to analyse differences between towers. The Bray-Curtis dissimilarity index was used to create a dissimilarity matrix and non-metric multidimensional scaling (NMDS) was used to visualize the data. The data points were then clustered according to the physical, chemical, and biological parameters. ANOSIM was used to test the statistical significance and strength of clustering correlation. The source of the treated water feeding the cooling towers was the only parameter that created significantly different clusters (Figure 5A) in agreement with hydrological basin (Figure 5B). The towers fed by the Ottawa river (located in Hull) and the ones fed by the St-François river (located in Sherbrooke) had the highest dissimilarity (pairwise test: R = 1, *P =* 0.005). The towers fed from the St-Lawrence river (located in Montreal, Monteregie, and Quebec) and the towers fed with a mixture of water from St-Charles Lake and the St-Lawrence river (Quebec) clustered together (pairwise test: R = −0.07, P = 0.7).

**Figure 5:**
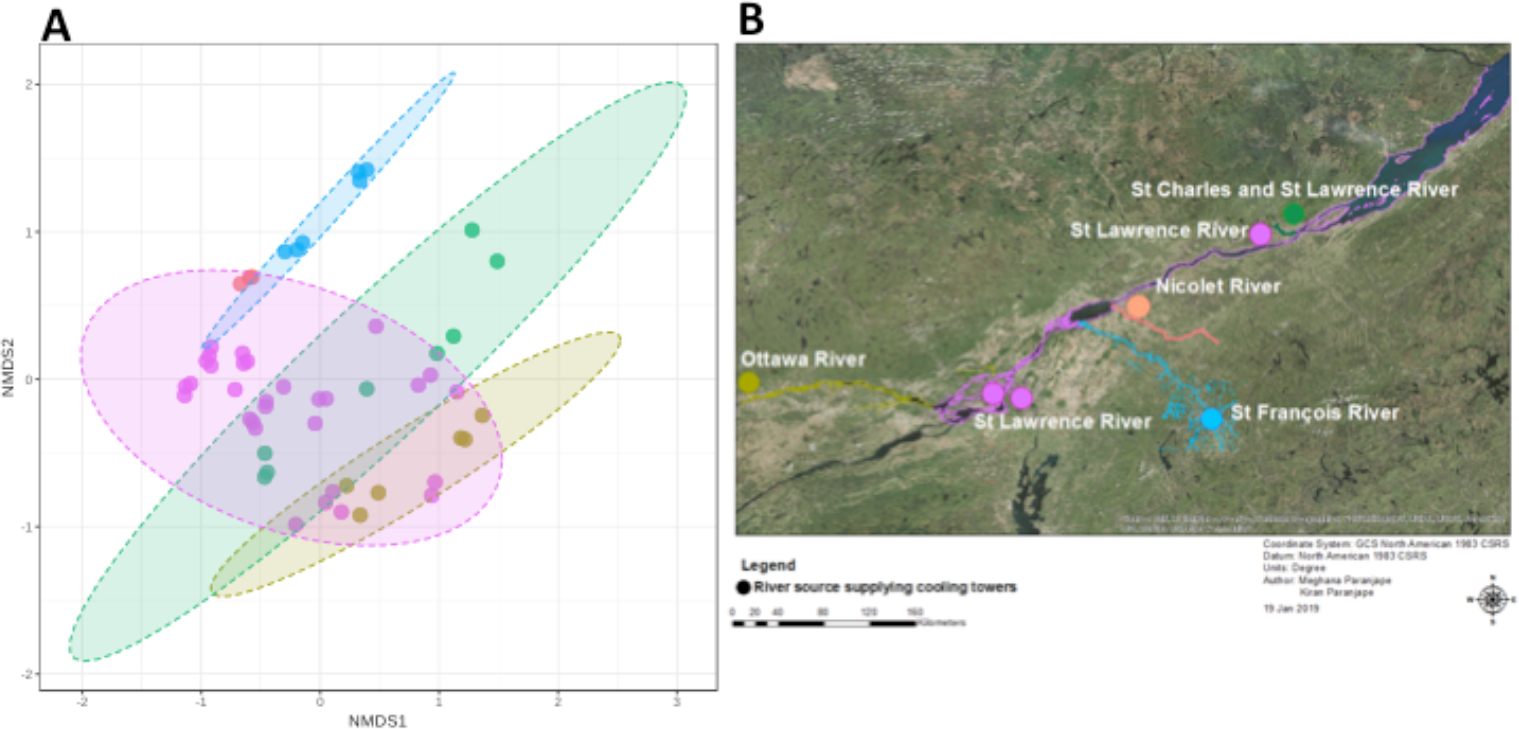
(A) Non-metric Multidimensional Scaling plot of tower microbiomes grouped by source of the water (stress = 0.1866). ANOSIM was used for statistical testing (R = 0.3927, P < 0.001). (B) Locations of the cooling towers sampled are indicated and colored according to the source of the water: Ottawa river (yellow), St-Lawrence river (pink), a mixture of water from St-Charles lake and St-Lawrence river (green), St-François river (blue), and Nicolet river (salmon).

### Correlation between the microbiome and key genera

Next, the prevalence of different microorganisms in the cooling towers was investigated to determine the core community of cooling towers. Seven out of the 72 genera showed prevalence above 80%, including *Pseudomonas*, *Porphyrobacter*, *Methylobacterium*, *Blastomonas*, and unclassified genera from the *Methylophilaceae*, the *Burkholderiaceae*, and the *Sphingomonadaceae* families (Supplementary Figure S4). *Pseudomonas* and *Methylobacterium* have near 100% prevalence in all towers at a relative abundance of 0.001; however, as relative abundance levels increased, prevalence decreased, indicating that these organisms are prevalent in most towers but at different abundance levels. These organisms likely constitute the core community of cooling towers. Six other genera had prevalence between 50% and 80%, including *Limnobacter*, *Obscuribacteriales*, *Sphingomonas*, *Sphingopyxis*, *Novosphingobium*, and *Bosea*. These organisms may be part of a transient community or may depend on specific physical and chemical parameters only found in a subset of cooling towers.

LEfSe was used to identify genera of importance for the different conditions studied. LEfSe is a machine-learning algorithm that uses a mix of statistical testing, linear discriminant analysis (LDA), and effect size to find the taxa that most likely explain the difference between specific parameters [59]. The algorithm was able to find significant taxa for most conditions; however, we decided to focus on the conditions where *Legionella*, *Pseudomonas*, or *Mycobacterium*, were distinguishing features. *Legionella* is enriched in conditions with low levels of total chlorine, medium levels of conductivity, and in towers with daily application of chlorine (Figure 6 and Supplementary Figure S5A). Conversely, *Pseudomonas* is enriched in towers with high levels of total chlorine, high levels of suspended solids, and with continuous application of chlorine (Figure 6 and Supplementary Figure S5B).

**Figure 6:**
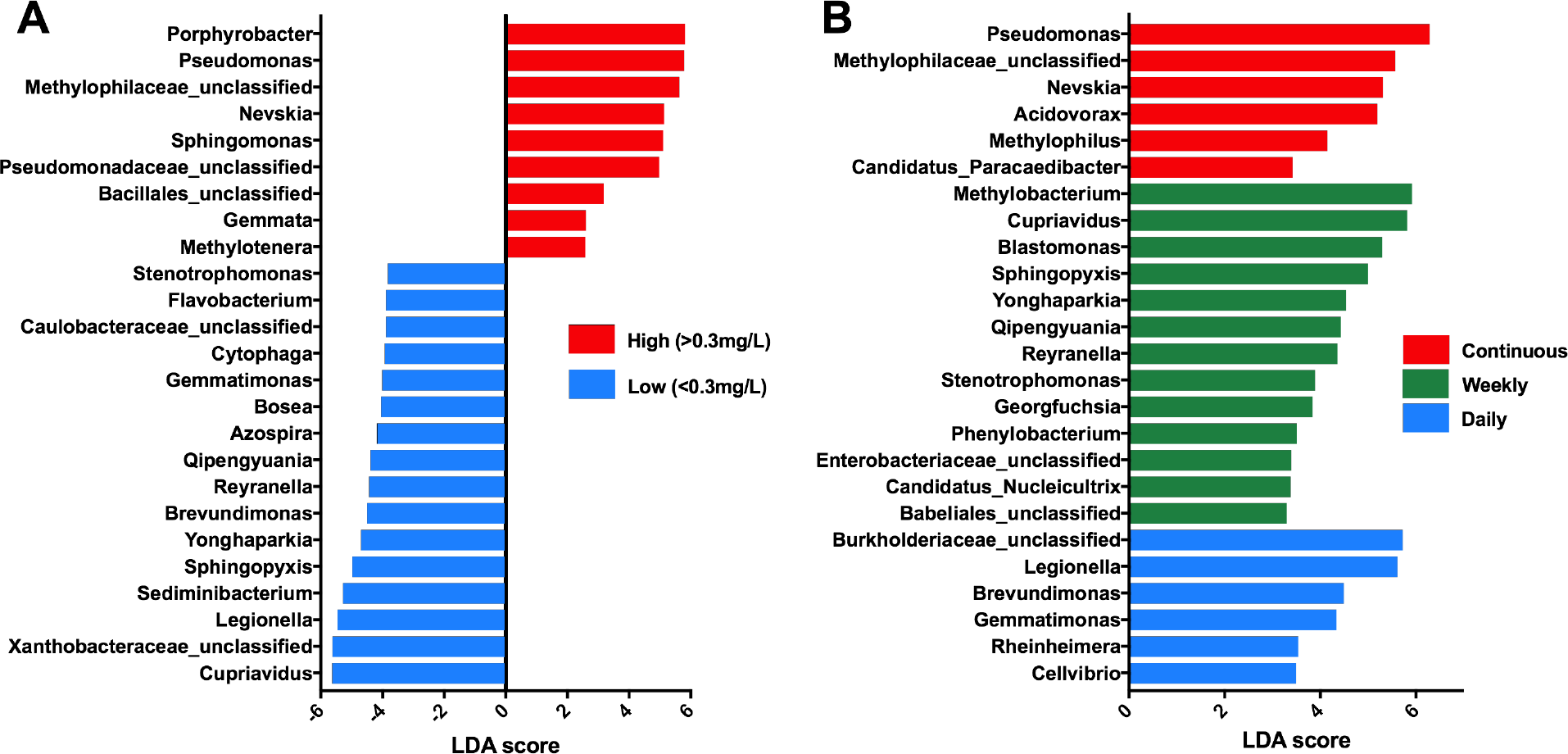
The machine learning algorithm LEfSe was used to identify significant taxa associated with chlorine concentrations (A), and with daily, weekly, and continuous application of chlorine (B). The LDA score is an effect size that measures the importance of the taxa in the condition studied.

LEfSe was then used to identify genera enriched in towers with *Legionella* and with *L. pneumophila* (Figure 7). Fifteen taxa were enriched in *Legionella*-positive towers and *Pseudomonas* was the only taxon enriched in towers without *Legionella* (Figure 7A). This analysis is in good agreement with a Spearman’s correlation analysis (Supplementary Figure S6). Several of the bacterial groups enriched in the *Legionella*-positive towers are unclassified or poorly studied, indicating a potential pool of uncharacterized interactions between *Legionella spp.* and these less well studied bacterial groups. Seven genera were enriched in *L. pneumophila*-positive towers (Figure 7B), including *Xanthobacteraceae* family, *Obscuribacterales* order, and the *Qipengyuania* genera. On the other hand, *Sphingobium* was the only genus enriched in towers that tested negative for *L. pneumophila* (LDA of 4.59).

**Figure 7:**
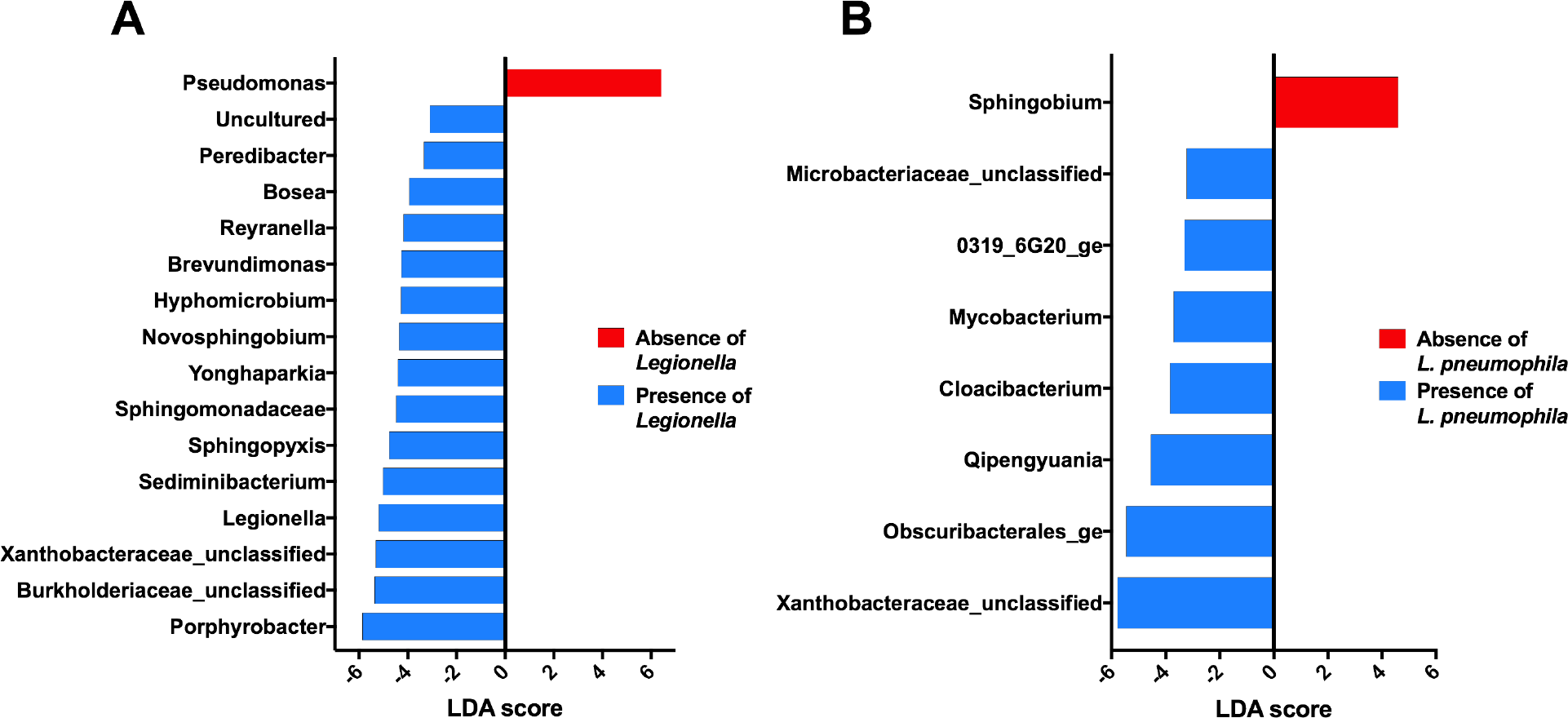
The machine learning algorithm LEfSe was used to identify taxa enriched in towers with and without *Legionella spp.* (A), and with towers with and without *L. pneumophila* (B). The LDA score is an effect size that measures the importance of the taxa in the condition studied.

## DISCUSSION

This study provides a snapshot of the ecology of the bacterial community of cooling towers in Southern Quebec. We hypothesized that the resident microbial population influences the colonization, survival, and proliferation of *L. pneumophila* in cooling towers. The source of the water was the main factor explaining the difference in the microbial composition of the cooling towers in our study (Figure 5A). The St-Francois River and the Ottawa River are distinct hydrological basin resulting in distinct microbiomes in the cooling towers respectively fed by these sources. The towers fed by the St-Lawrence river showed similar microbiomes and overlapped with the towers fed with a mixture of water from St-Lawrence river and the St-Charles lake (CN 2, 3 and 4). The Ottawa River feeds into the northern shore of the St-Lawrence river at the west of Montreal and the St-François river is a tributary joining the St-Lawrence river about 160 km downstream of Montreal, (Figure 5B). Both rivers probably have minimal impact on the St-Lawrence river microbiome. Taken together, our results suggest that the microbial composition of the source water dictates the microbial population of the cooling towers; however, other parameters associated with geographic location are likely to play a role. For example, the airborne microbiome could be a confounding factor. Other parameters did not create significantly different clusters. Although chlorination schedule clearly affects the microbial diversity in cooling towers (Figure 3), its effect is non-specific.

Generally, the bacterial community were dominated by species from the *Proteobacteria* phylum. This is in agreement with several other studies that looked at cooling towers and other EWS [11, 50, 60, 61, 62, 63, 64]. While *Actinobacteria* and *Proteobacteria* dominate in equal proportions freshwater sources feeding EWS, the *Actinobacteria* population is greatly and significantly reduced in EWS, leaving the *Proteobacteria* as the dominant phylum of these environments [65, 66, 67, 68, 50, 61]. Water treatment increases levels of certain groups of *Alphaproteobacteria*, such as *Sphingomonadaceae*, *Beijerinckiaceae*, and *Rhizobiaceae* [69]. Stagnation of water in pipes also contributes to increase levels of *Proteobacteria* [61]. Since all cooling towers in our study are fed with treated municipal water, the dominance of *Proteobacteria* was expected.

Some genera present in the cooling towers are frequently observed in other EWS, whereas others are less frequently identified. *Pseudomonas*, *Blastomonas*, *Methylobacterium*, and unclassified genera from the *Bukholderiaceae* family constitue the core microbiome of cooling towers. The high prevalence of *Methylobacterium* indicates that methylotrophy could be an important ecological function in cooling tower. *Limnobacter*, *Sphingopyxis*, *Novosphingobium*, *Bosea* were only found between 50 to 60% of towers (Supplementary Figure S4). Our results differ somewhat compared to other studies. For instance, a two-year study of a German cooling tower showed high abundance of the environmental *Proteobacteria* ARKICE-90, *Nevskia* genus, *Methilophilus*, and uncultured bacteria from the family *Cytophagaceae*, but relatively low abundance of *Pseudomonadales* and absence of *Methylobacterium* [50]. Another study that looked at cooling towers of pharmaceutical plants and oil refinery in Italy and Eastern Europe also found high levels of *Proteobacteria*, such as *Rhodobacteraceae*, *Sphingomonadaceae*, *Bradyrhizobiaceae*, as well as *Cyanobacteria*, but no *Pseudomonas* [70]. Thus, the community composition seems influenced by the intrinsic properties of a cooling tower and its geographic location. For instance, piping material, disinfection strategies, water sources, nitrate concentrations, iron concentrations, water treatment, dissolved organic carbon, and seasonality are all factors that have been shown to shape the bacterial population of different EWS [71, 72, 73, 74, 75, 76, 61].

In our case, several physico-chemical parameters affected the microbiome of cooling towers (Supplementary Figure S2). *Legionella* was enriched in towers with low levels of total chlorine (<0.3mg/L) and with daily applications of chlorine whereas *Pseudomonas* was enriched in towers with high levels of chlorine and continuous application (Figure 6). These findings suggest that continuous application and maintenance of a free chlorine residual greater than 0.3 mg/L is key to prevent the colonization of cooling towers by *Legionella*. From the data, three possible mechanisms may explain this phenomenon. First, the most obvious explanation is that these parameters ensure sufficient concentration and contact time to inactivate *Legionella* [77]. The second explanation is linked to the decrease in alpha diversity caused by a continuous application of chlorine (Figure 3C), potentially restricting the growth of species beneficial for *Legionella spp*. The continuous presence of residual chlorine reduces the concentrations and diversity of host cells and the biofilm mass [78, 24], thus limiting the possibility for increased resistance of *L. pneumophila* through integration into the biofilm and inside protozoan hosts [79, 19]. As seen in Figure 7A, *Legionella* was positively correlated with many genera, which could promote its survival and proliferation. For instance, *Reyranella*, *Brevundimonas*, *Sphingopyxis*, and *Yonghparkia* are enriched in *Legionella-*postitive towers with low level of chlorine and treated by periodic application. These genera may either directly or indirectly promote the growth of *Legionella spp*. Alternatively, these taxa may be indicators of environmental condition permissive for the presence of *Legionella*. The third possible explanation for the lack of *Legionella spp.* in towers with high levels of chlorine and continuous application may be linked with the presence of *Pseudomonas spp.* in these towers. This was demonstrated by using LEfSe and Spearman’s correlation, which showed that these two genera were the most negatively correlated to one another (Figure 7A and Supplementary Figure S6). In contrast, the relation between the genus *Pseudomonas* and the species *L. pneumophila*, detected by qPCR, was less clear, since *Pseudomonas* is not a significant taxon in towers without *L. pneumophila* (Figure 7B). However, the average number of *Pseudomonas* reads were significantly lower (Mann-Whitney, *P <* 0.05) in *L. pneumophila* positive towers (920 reads) than in negative towers (5476 reads). Similarly, *Llewenlyn et al.* reported higher abundance of *Pseudomonadaceae* in *Legionella*-negative towers [11]. Thus, continuous chlorination promotes the establishment of a *Pseudomonas* community, lower alpha diversity, and low levels of *Legionella spp*. A positive correlation between chlorine and *Pseudomonas* was previously reported [72, 74, 80]. *P. aeruginosa* has a higher tolerance to chlorine than other water-borne bacteria, which is attributed in part to biofilm formation [81, 82, 83, 84, 85]. On the other hand, many species of *Pseudomonas* are highly competitive and possess many mechanisms to outcompete other bacteria, such as type VI secretion systems, pyoverdine, phenazine, and metabolic flexibility [86, 87, 88, 89, 90]. The fact that *Pseudomonas spp*. is negatively correlated with alpha diversity further adds evidence to the competitive nature of *Pseudomonas* (Figure 4C). Many species of *Pseudomonas* inhibit the growth of *L. pneumophila* on CYE agar by producing antagonistic diffusible compound [91, 92]. Furthermore, *L. pneumophila* is unable to persist in biofilm produced by *P. aeruginosa* [27]. Consequently, *Pseudomonas spp.* may directly restrict the presence and growth of *Legionella spp* in water system. In addition, *Pseudomonas* could act on *Legionella spp.* indirectly. *P. aeruginosa* is known to kill the amoeba *A. castellanii*, a host cell of *L. pneumophila*, by secreting toxic effector proteins using the type III secretion system [93] and could therefore reduce the pool of host cells. *Pseudomonas* may also inhibit the growth of certain bacterial species that promote the growth of *Legionella spp.* or that are preys for host cells. Thus, the data suggest that high concentration of chlorine applied continuously inhibit the colonization and proliferation of *Legionella spp.* but promote the establishment of a *Pseudomonas* community. This may be of concern for tower maintenance, as *P. aeruginosa* is an opportunistic pathogen of great concern [94].

Finally, our results seemed to indicate that *Legionella spp.* and *L. pneumophila* are associated with several other genera. Spearman’s correlation and LEfSe analysis showed that several taxa were positively correlated and enriched in towers containing a population of *Legionella* (Figure 7 and Supplementary Figure S6). Of note, the family *Xanthobacteraceae* was positively correlated with both *Legionella spp.* and *L. pneumophila*. Many members of this family are chemolithoautotrophs and some are able to fix nitrogen [95]. Therefore, they are likely at the bottom of the food chain and could feed *L. pneumophila* host cells. In addition, several isolates are able to degrade chlorinated and brominated compounds (*Oren, 2014*). An uncultured *Xanthobacteraceae* was recently identified as a component of biofilm growing in a model hot water system colonized by *L. pneumophila* [33]. It is tempting to speculate that *Xanthobacteraceae* could help the development of healthy biofilms by producing organic molecules and reducing local concentration of disinfectant or toxic by-product, which in turn could promote *L. pneumophila* colonization. The genus *Sphingobium* was the only one negatively correlated with *L. pneumophila*. Species from this genus may be associated with free-living amoeba [96]. Although it is not clear if this genus contains species that can grow within amoeba, it can be hypothesized that *Sphingobium* could compete with *L. pneumophila* for host cells, which would result in lower *L. pneumophila* growth. So far and to the best of our knowledge, none of these taxa have been documented to interact with *Legionella* species. Furthermore, several of these taxa are unclassified or uncultured organisms and thus their life cycle and ecological interactions are poorly understood. Potentially, the interaction of these different taxa and *Legionella* could be indirect as they may be prey for host cells. This would support the hypothesis that *Legionella* colonization of towers depends on the establishment of bacterial community that feeds the host cell population. Our findings support the notion that Legionnaires’ disease outbreaks may depend on a network of uncharacterized microbial interactions between *L. pneumophila* and the bacterial community, along with an optimal range of physical and chemical parameters, that promote its colonization, survival, and proliferation in cooling towers.

## CONCLUSION

In conclusion, three main observations emerge from this work. First, the source of the water is the main factors affecting the bacterial community of cooling towers. Secondly, the *Legionella* population itself is severely affected by the alpha diversity, the level of *Pseudomonas*, levels of chlorine, and most importantly, the frequency of chlorine treatment. Finally, our results indicate that *Legionella* and *L. pneumophila* could interact with several uncultured and unclassified taxa suggesting that colonization of towers and likelihood of outbreaks could be potentiated by as of yet uncharacterized interactions between *L. pneumophila* and several bacterial species. Therefore, it seems that the presence of *Legionella* in cooling towers is influenced by several factors that can be targeted to reduce the risk of outbreaks. In particular, continuous chlorine treatment seems to promote conditions associated with the absence of *Legionella*.

## ACKNOWLEDGMENTS

We are indebted to SQI for access to cooling towers and help with sampling. This work was supported by a FRQNT Team Grant to MP and SPF. We thank Meghana Paranjape for helping us create the sampling map.

