## Supplemental Figures for "Presence of *Legionella* spp. in cooling towers: the role of microbial diversity, *Pseudomonas*, and continuous chlorine application"

[1]Department of Natural Resource Sciences, Faculty of Agricultural and Environmental Sciences, McGill University, Sainte-Anne-de-Bellevue, QC, Canada; [2]Department of Civil Engineering, Polytechnique Montréal, Montréal, QC, Canada; [3]Department of Food Science and Agricultural Chemistry, Faculty of Agricultural and Environmental Sciences, McGill University, Sainte-Anne-de-Bellevue, QC, Canada. [4]Department of Animal Science, Faculty of Agricultural and Environmental Sciences, McGill University, Sainte-Anne-de-Bellevue, QC, Canada.

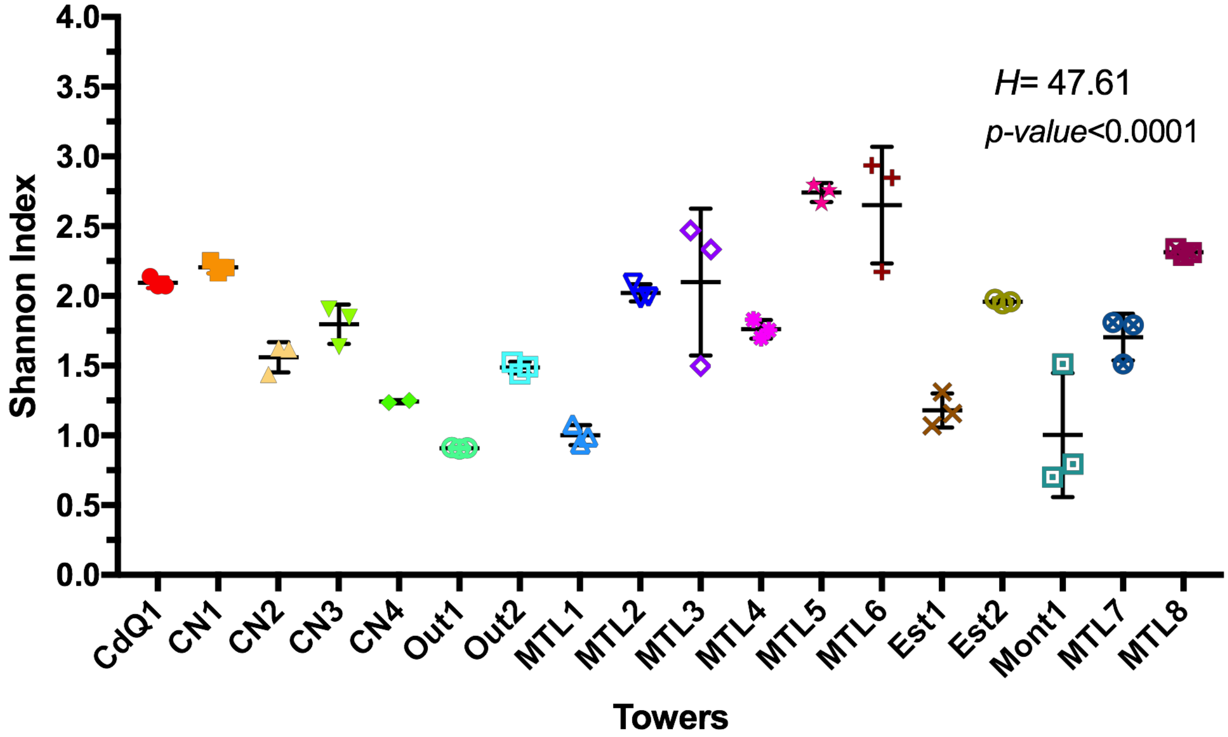


**Figure S1:** Alpha diversity levels of sampled cooling towers using the Shannon index. The Kruskal-Wallis test was used to assess statistical significance.

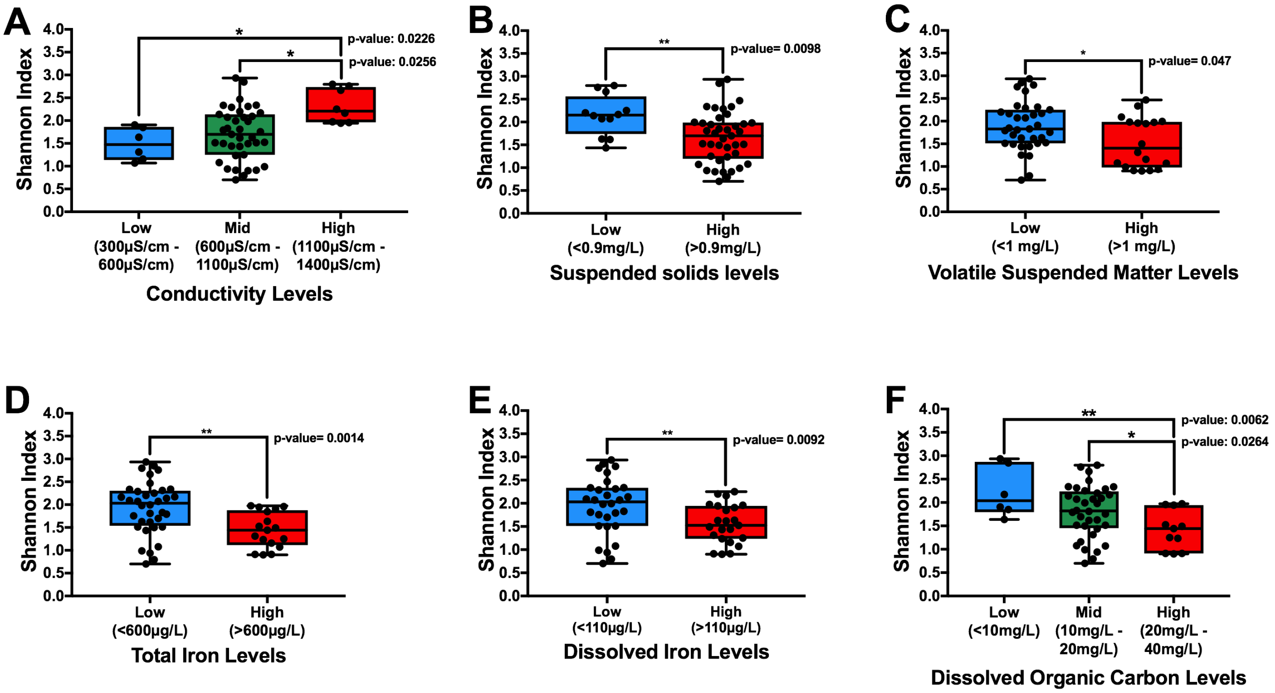


**Figure S2:** Effect of levels of conductivity (A), suspended solids (B), volatile suspended matter (C), total iron (D), dissolved iron (E), and dissolved organic carbon (F) on alpha diversity (Shannon index) of cooling towers. Higher levels of suspended solids, volatile suspended matter, total iron, dissolved iron, and dissolved organic carbon decreased alpha diversity. Higher levels of conductivity increased alpha diversity of towers.

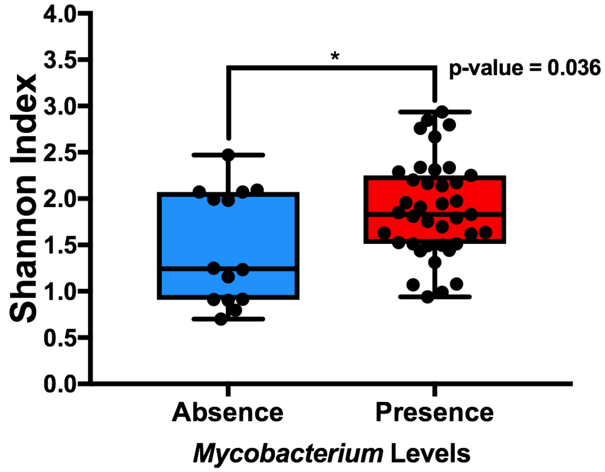


**Figure S3:** Alpha diversity and the presence of *Mycobacterium* in sampled cooling towers. Towers with the presence of *Mycobacterium* had higher levels of alpha diversity than towers without Mycobacterium.

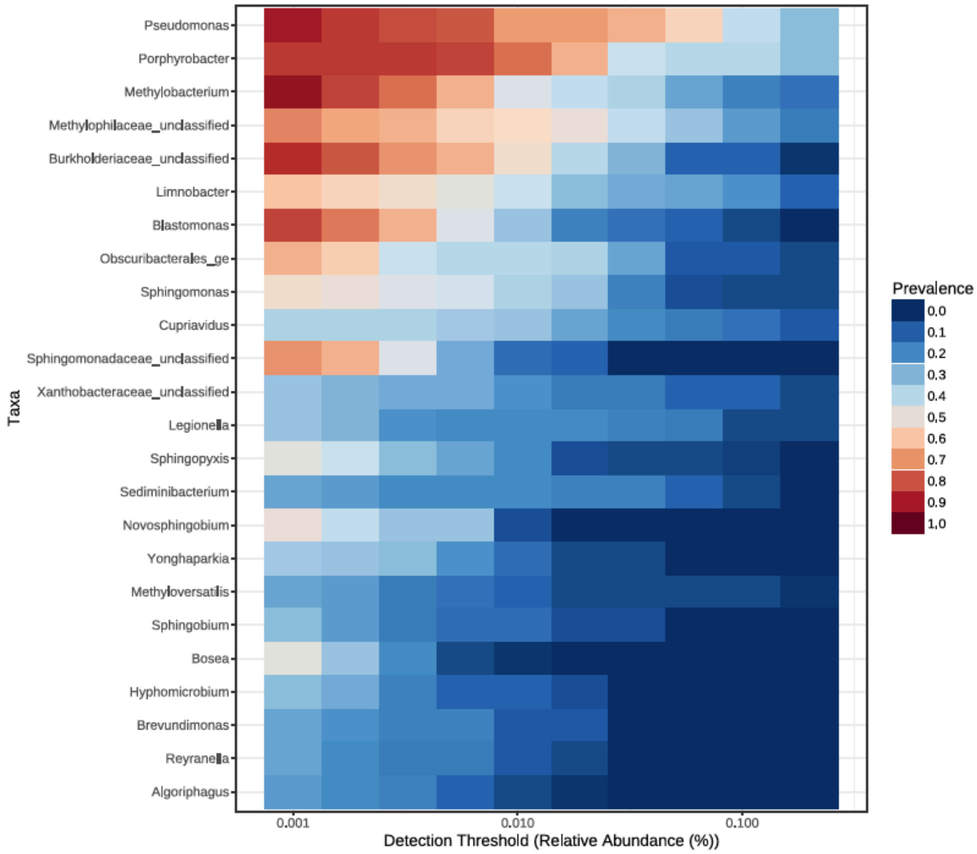


**Figure S4:** Most prevalent genera present in sampled cooling towers representing the core bacterial community.

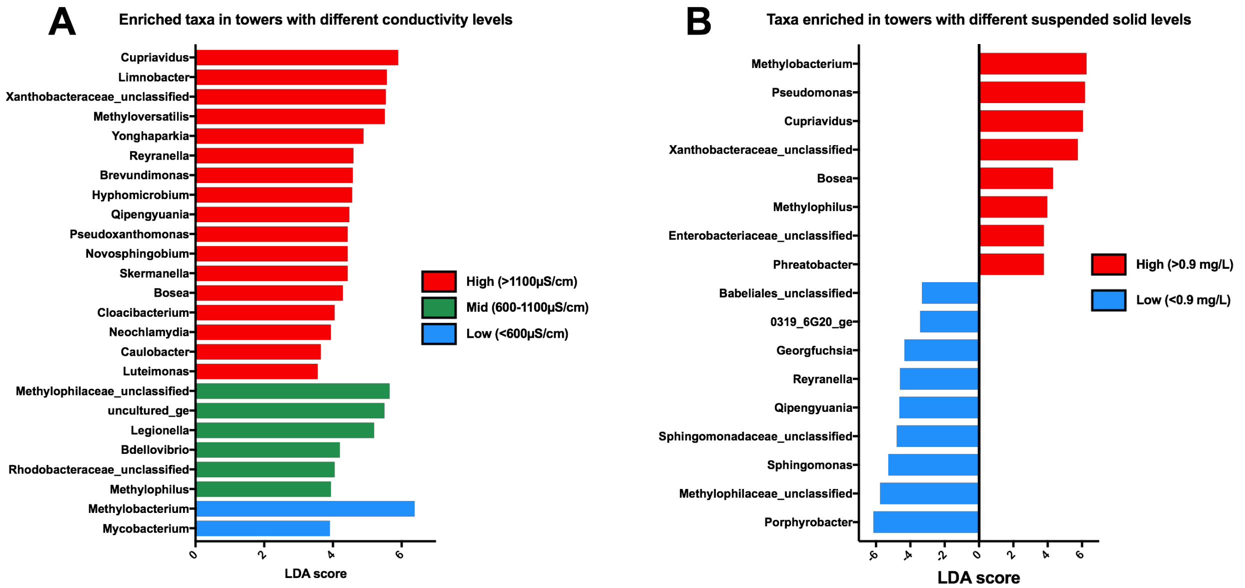


**Figure S5:** Enriched taxa in towers with different levels of conductivity (A), and with different levels of suspended solids (B) using LEfSe. *Legionella* was enriched in towers with mid levels of conductivity, and *Pseudomonas* was enriched in towers with high levels of suspended solids.

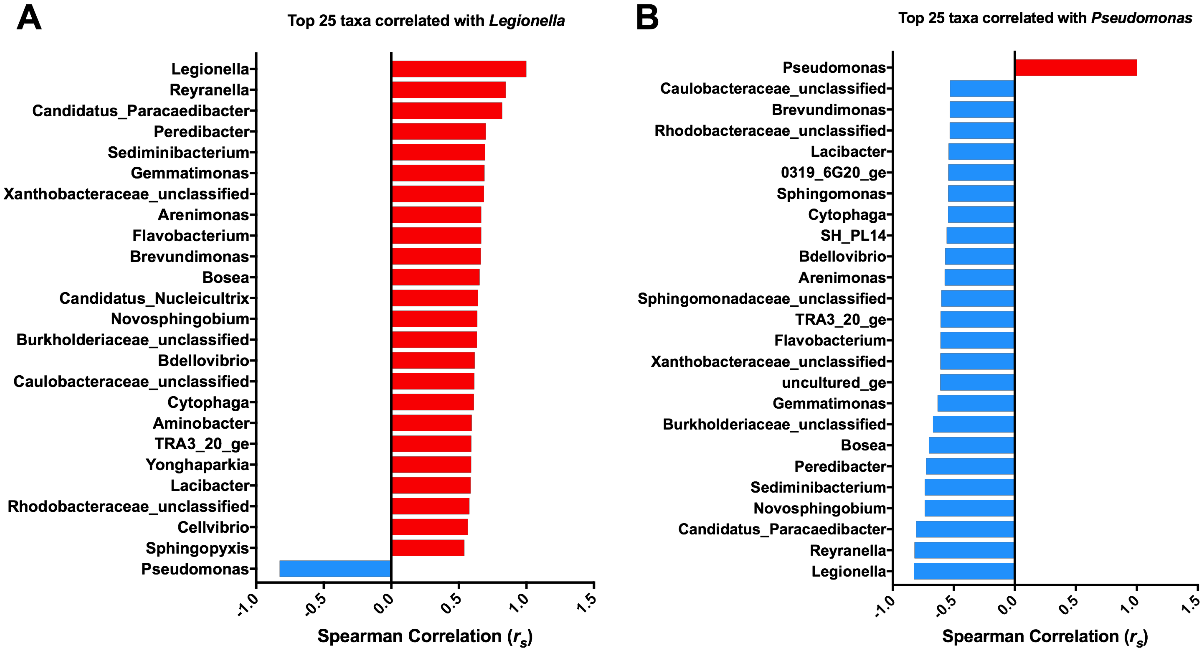


**Figure S6:** Spearman correlation of top 25 taxa correlated with *Legionella* (A), and with *Pseudomonas* (B). *Legionella* was positively correlated with many taxa; *Pseudomonas* was negatively correlated with most taxa.
